# Chromosome-level genome assembly of the Brazilian merganser (*Mergus octosetaceus*), a rare and elusive waterfowl species

**DOI:** 10.1101/2025.09.16.676261

**Authors:** Henry Paul Granger-Neto, Isaac Rafael Freitas Borges, Davi Barbalho Cavalcanti, Davidson P. Campos, Kathleen Horan, Jennifer Balacco, Brian O’Toole, Tatiana Tilley, Nivesh Jain, Linelle Abueg, Nadolina Brajuka, Giulio Formenti, Olivier Fedrigo, Erich D. Jarvis, Fabrício R. Santos

**Affiliations:** Departamento de Genética, Ecologia e Evolução, Universidade Federal de Minas Gerais, Minas Gerais, Brazil; Vertebrate Genome Laboratory, The Rockefeller University, New York, USA

**Keywords:** Brazilian merganser, *Mergus octosetaceus*, Chromosome-level genome, Reference genome

## Abstract

**Background:** The Brazilian merganser (*Mergus octosetaceus*) is a critically endangered bird, with fewer than 250 mature individuals remaining in fragmented populations of the Cerrado biome in Brazil. It is an ecologically demanding species that requires clear water of fast-flowing rivers for diving and foraging, and riverside cavities for nesting. Technical and logistical challenges have restricted sampling and research to few genetic studies so far. The genetic results consistently revealed a low genetic diversity and high inbreeding in the species. Besides, the lack of a reference genome has precluded deeper evolutionary and conservation genetic analyses for the species.

**Results:** We generated the first high-quality, chromosome-level genome assembly for *Mergus octosetaceus* using PacBio HiFi long reads and Arima Hi-C data from a female specimen. The assembly is 1.25 Gb in length, is highly complete, scoring 98.9% of BUSCO completeness, and includes both sex chromosomes (Z and W). Annotated repetitive elements comprise 18.94% of the genome, with evidence of recent activity, particularly among LINEs and LTRs retrotransposons. We also produced and annotated the complete mitochondrial genome, identifying 37 genes and the control region. Comparative synteny analysis with other Anatidae species revealed strong chromosomal conservation, with several inversion events in macrochromosomes. Demographic history reconstruction indicated fluctuations in effective population size, with significant reductions overlapped by major temperature changes, highlighting potential climate sensitivity of this lineage.

**Conclusion:** This reference genome provides a fundamental resource for *M. octosetaceus*, enabling insights into genome evolution, chromosomal dynamics within Anatidae, and past population history. It is a remarkable foundation for future research and conservation strategies for managing and restoring this critically endangered species.

## Introduction

The Brazilian merganser (*Mergus octosetaceus* Vieillot, 1807) has been classified as critically endangered for over 30 years [1] and ranks among the world’s rarest waterfowl species. Since the extinction of *Mergus australis* in the early 20th century, it remains the sole representative of the tribe Mergini in the Southern Hemisphere [2].

Historically, its distribution included the Brazilian Cerrado and Atlantic Forest, as well as bordering parts of Paraguay and Argentina [3], where it has been considered locally extinct for at least 23 years [4–6]. Current populations are restricted to four geographic groups that were divided into three genetic management subpopulations [7], all located in the Brazilian Cerrado. The wild population is estimated to be fewer than

250 mature individuals with a declining trend [8]. Besides, since 2011 a captive population has been established for the species [2], which is now composed of about 50 individuals descended from founders of all four geographic locations in Brazil, derived from collected eggs in wild nests.

Brazilian mergansers exhibit extreme sensibility to human disturbance due to specific ecological requirements: they depend on clear, fast flow waters for diving-based foraging and cavities for nesting, including hollow tree trunks or cavities in the rocks or riverbanks [9]. Breeding pairs are territorial, often abandoning large river stretches following contact with humans, and conflicts among individuals are common.

Like many threatened species, genetic studies face challenges due to limited accessibility, resources, and logistical constraints in sample collection. Consequently, research has relied on partial genomic approaches (e.g., mtDNA, microsatellites, GBS, ddRAD) as alternatives in the absence of a reference genome.

Only seven genetic studies were performed with the species [7,10–15], all indicating low genetic diversity, high kinship, and inbreeding in both wild and captive populations. Six of those studies originated from our group and, notably, the last two [7,14] have used reduced representation genomic libraries (GBS and ddRAD) to identify SNPs with a de novo approach (without a reference genome).

A high-quality reference genome enables precise genetic inference, increases the robustness of studies, and enhances efforts for the preservation of the genetic heritage [16]. Advances in long-read sequencing, assembly algorithms, and the invaluable work of consortia like the Vertebrate Genomes Project (VGP), European Reference Genome Atlas, Darwin Tree of Life, Genomics of the Brazilian Biodiversity, Genotropics and Earth BioGenome Project now make this achievable [16–21].

Through collaboration with VGP, and integrating PacBio HiFi and Hi-C data, we present the first chromosome-level genome assembly for *Mergus octosetaceus*. Preliminary analyses reveal a slight increase in repetitive DNA, possibly due to better assembly/annotation, and high syntenic conservation when compared to other Anatidae, while the ancestral demographic pattern seems to be responsive to past climatic fluctuations. We expect this reference genome will support future evolutionary research and conservation management strategies for the species and provide increased power for comparative studies within the Anatidae.

## Methods

### Sampling and permits

A fresh blood sample (5 ml) was obtained from a captivity individual (Studbook #49-AAA001735/Mo242) maintained at Itatiba Zoopark (São Paulo State, Brazil) according to the ethical protocol 239/2018 (CEUA-UFMG), as part of the reproductive management program of the species according to the National Action Plan of Conservation of *Mergus octosetaceus* (Portaria 44, April/08/2014, Ministry of Environment of Brazil). The access to genetic resources was registered at SisGen A324339 of the Brazilian system of genetic heritage. A tissue voucher (CCT_AV2000387) of the sequenced individual is stored at the Centro de Coleções Taxonômicas of Universidade Federal de Minas Gerais.

### Genome Sequencing and Assembly

The genome was sequenced and assembled using the VGP v2.0 pipeline in the Galaxy workflow space [22]. In brief, genomic DNA was extracted from 10 µl of blood and purified using a Bionano SP DNA kit (PN 80042). DNA quantity was measured using a Qubit 3 fluorometer dsDNA Broad Range Assay (Invitrogen cat. no. Q32850). DNA sizing was assessed with an Agilent Femto Pulse. The DNA was fragmented to ~20-40 Kb range, using a Megaruptor 3 (Diagenode, Denville, NJ, USA), speed 30, and standard hydropores (Cat. No. E07010003). Then 5 µg of the sheared DNA was used to prepare a PacBio HiFi library using SMRTbell prep kit 3.0 (Pacific Biosciences PN 102-182-700). The final HiFi library was size selected to remove inserts under 10 kb using a PippinHT instrument (Sage Science, Beverly, MA, USA). The HiFi library was sequenced on 8M SMRT cells and a Sequel IIe instrument using Sequel II sequencing kit 2.0 (101-820-200), 2-hour pre-extensions, and 40-hour movie time.

For Hi-C sequencing, 50 µL of blood was used to prepare an Arima Hi-C library with an Arima-HiC 2.0 kit (Arima Genomics, Carlsbad, CA, USA) following the manufacturer’s protocol. The Hi-C library was sequenced with 2×150 bp read length with the Illumina NovaSeq platform.

For genome assembly, the Hifiasm v0.19.3 in Hi-C integrated mode was used to assemble haplotyped phased contigs (haplotype 1 and haplotype 2 representing chromosomal segments of each parent). Each haplotype was then separately scaffolded into chromosomes, using the same Hi-C data and the yahs v1.2a.2 algorithm. Manual curation was performed in dual curation mode with both haplotypes, to fix structural errors and name chromosomes.

### Genome Curation

Manual genome curation was primarily performed using PretextView [23], and supplemented by JBrowse2 [24] and HiGlass [25]. Scaffold integrity was assessed and refined according to the methodology of Howe et al. [26], which included resolving contamination, correcting mis-joins and missed joins, and removing flagged duplicate sequences. The curation workflow, including quality control steps and decision criteria, is publicly archived at https://gitlab.com/wtsi-grit/rapid-curation (manuscript in preparation).

### Genome Statistics

For both primary and curated assemblies, the genome was evaluated using gfastats v1.3.9 [27] for assembly statistics and BUSCO v5.8.2 [28] with the lineage aves_odb10 for completeness. The snail plot summarizing major statistics was generated with BlobToolKit v4.4.0 [29].

### Repeat Annotation

Repeat annotation was performed using the EarlGrey v5.1.1 pipeline [30]. The resulting library was first automatically curated by TEtrimmer v1.4.0 [31], and further manually curated with TE-Aid v.1.1 [32] to evaluate consensus sequence divergence, genome coverage, and TE protein annotations. Structural features of repeats, including tandem repeat localization, were analyzed in Geneious v8.0.5 [33] using dot plots and genome mapping. Functional annotation of TE proteins was conducted using InterProScan v5.73-104.0 [34]. Finally, the repeat content was quantified with RepeatMasker v4.1.6 [35] using default settings to generate the genome mask.

### Mitochondrial annotation

We annotated the mitochondrial genome through an integrated approach, starting with gene identification using MITOS2 [36] followed by precise control region (CR/D-loop) delineation using MitoAnnotator [37] and coding and intronic regions validation with MFannot [38]. Finally, we compared the final annotation against a museum-specimen mitogenome of *M. octosetaceus* [15] to assess the structural conservation of the annotated features.

### Collinearity analysis

For the evaluation of collinearity conservation, we decided to use only chromosome-level genomes from the Anatidae family. One representative of each tribe within the Anatinae subfamily was selected: *Anas platyrhynchos* (Anatini), *Aythya fuligula* (Aythini), *M. octosetaceus* (Mergini), *Alopochen aegyptiaca* (Tadornini), and *Cairina moschata* (Cairinini). Additionally, we included representatives from the subfamilies Anserinae (*Cygnus atratus*) and Oxyurinae (*Oxyura jamaicensis*), with *Gallus gallus* as the outgroup.

Genomes were filtered using SeqKit v2.8.2 [39] to retain only autosomes and the Z chromosome. Scaffolds, unlocalized/unplaced sequences, and the W chromosome were excluded. Pairwise whole-genome alignments were performed with minimap2 v2.28 [40] for cross-species comparisons. Chromosomal syntenic relationships were visualized using NGenomeSyn v1.41 [41].

### Demographic history reconstruction

In brief, HiFi reads were mapped to the reference genome using minimap2, followed by BAM file sorting and indexing with samtools v1.20 [42]. Using GATK v4.5.0.0 [43], duplicates were marked, haplotypes were called, and genotypes were generated. The resulting BCF file was filtered with bcftools v1.20 [44], masked using repetitive region coordinates from a BED file, and used to generate a diploid consensus sequence. SeqKit then removed sexual chromosomes, chromosomes smaller than 5 Mbp, and all scaffolds.

Historical effective population size (N_e_) was inferred using Pairwise Sequentially Markovian Coalescent (PSMC) v0.6.5 [45] with a coalescent window of 1+1+1+1+25*2+4+6, a generation time of 2 years, and a mutation rate of 5 × 10^−9^ (parameters tested as described in **Supplementary Fig. S1–S2**).

## Results

### Genome assembly and quality control

We generated 56.8 Gb of HiFi long-read data (~46.2× coverage) with a mean insert size of 17 kb and 133.4 Gb of Arima Hi-C data (~115.15× coverage) from a female specimen from Serra da Canastra region. Using the HiFi sequencing for the primary assembly and the Hi-C data for curation and polishing, we produced the first high-quality chromosome-level genome for *M. octosetaceus*.

The final assembly comprises two haplotypes: bMerOct1.hap1 (haplotype 1) and bMerOct1.hap2 (haplotype 2). Key assembly statistics are summarized in **Fig. 1A** (haplotype 1) and **Fig. 1C** (haplotype 2), with full metrics in **Supplementary Tables S1– S2**. Haplotype 1 was the most complete assembly, with a total length of 1.25 Gbp, 98.9%

**Figure 1.**
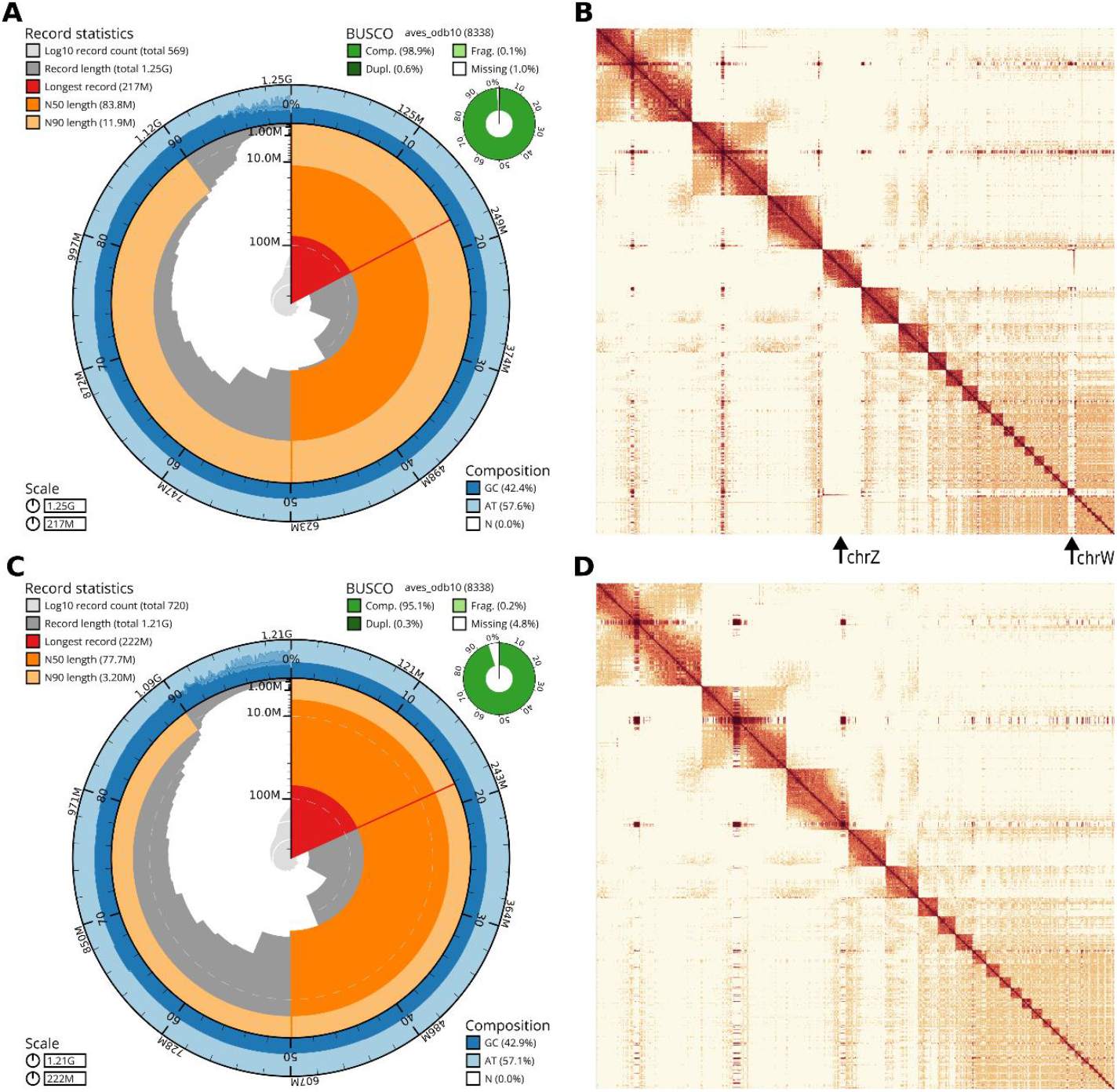
Haplotype assembly evaluation and curated Hi-C chromosome mapping. **A –** Snail plot for the haplotype 1. **B** – Curated Hi-C reads mapping for each chromosome of haplotype 1. **C –** Snail plot for the haplotype 2. **D –** Curated Hi-C reads mapping for each chromosome of haplotype 2.

BUSCO completeness, an N50 of 83.8 Mbp, and included both sex chromosomes (Z and W). Hi-C contact maps for the 26 largest chromosomes are presented in **Fig. 1B** (haplotype 1) and **Fig. 1D** (haplotype 2), showcasing robust chromosome-level support.

### Repeat Annotation

We identified that 18.94% of the genome of the Brazilian merganser consists of repetitive elements (**Fig. 2A**), a proportion consistent with typical avian genomes [46] but notably higher than those of its close relatives, such as the chicken (*Gallus gallus*) and mallard (*Anas platyrhynchos*), as reported in RepeatMasker [47]. Many of the elements are “recent”, especially long interspersed nuclear elements (LINEs) and long terminal repeats (LTRs), as indicated by Kimura’s divergence values (**Fig. 2B**). Finally, **Fig. 2C** illustrates the genome-wide distribution of repetitive sequences, with blue peaks indicating centromeric regions **–** linked to the accumulation of satellite repeats in Eukaryotes **–** highlighting their enrichment in these structural domains.

**Figure 2.**
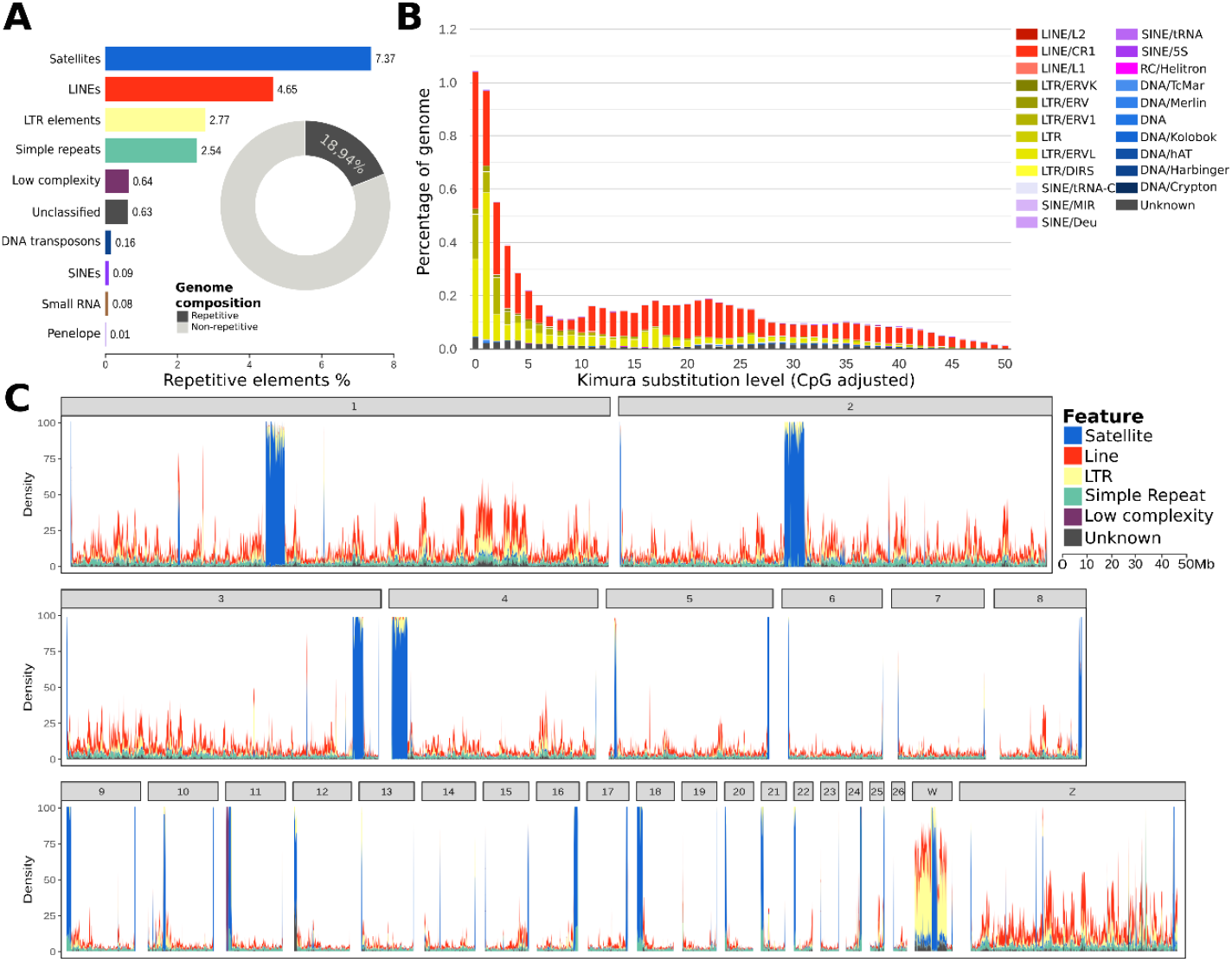
Repetitive genome annotation. **A** – Proportion of major repetitive element classes in the genome. **B** – Kimura Two-Parameter (K2P) distribution of repetitive elements. **C** – Major classes repetitive elements distribution across main chromosomes (i.e. >5 Mpb).

### Mitogenome assembly and annotation

The complete mitochondrial genome of *M. octosetaceus* (**Fig. 3**) is 16,621 bp in length, of which 68.60% represent coding regions, and has a GC content of 48.52%. As expected for vertebrates, we found 37 complete mitochondrial genes: 22 transfer RNAs (tRNA), 13 protein-coding genes (CDS), and two ribosomal RNAs (rRNA). The ND3 gene is divided into two subunits, as found in other Anatidae [48]. Additionally, the replication control region (CR/D-loop) was identified between the tRNA-TTC and tRNA-GAA genes, but only the heavy strand replication origin (OH) was annotated (expected for birds).

**Figure 3.**
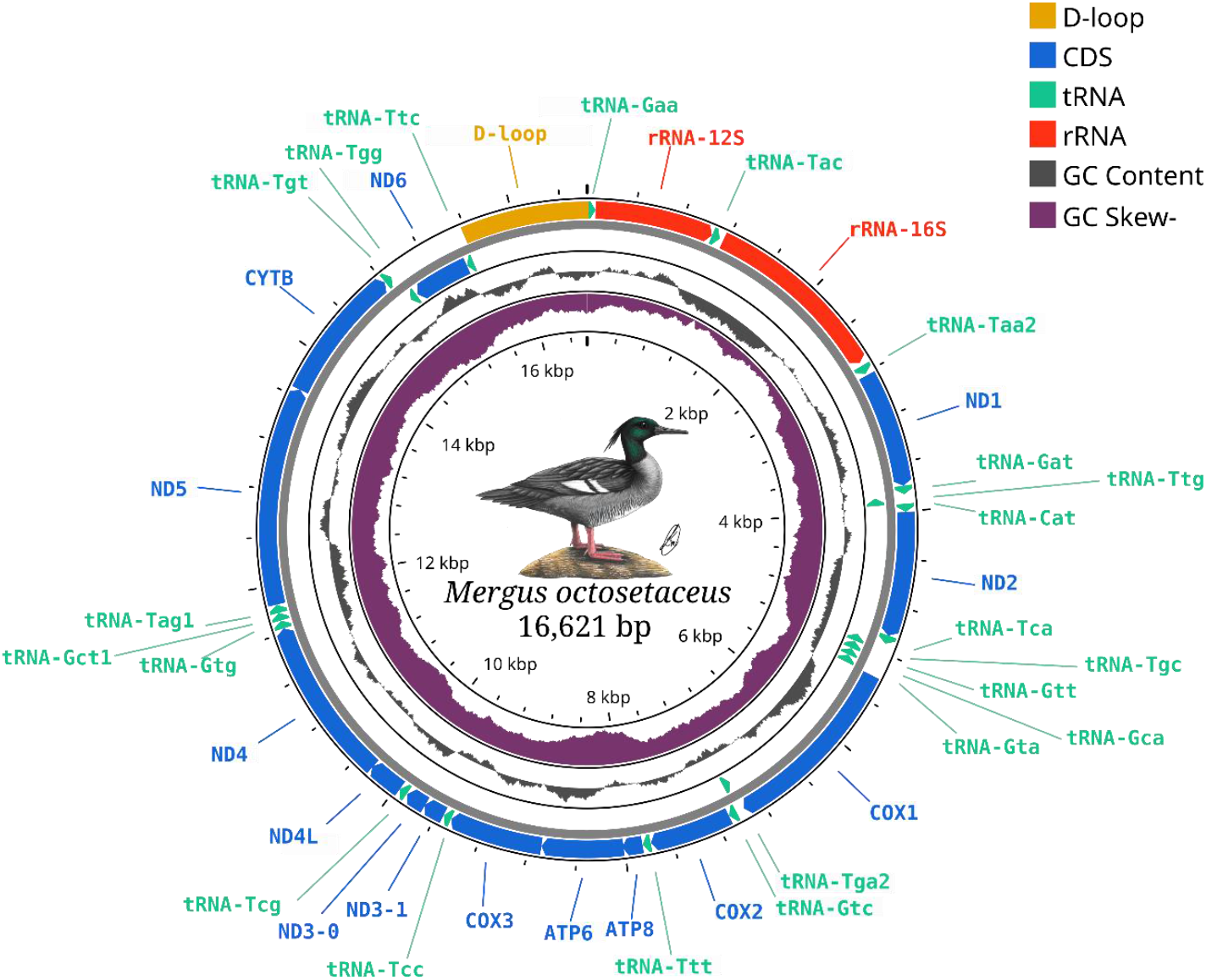
Structural annotation of the *Mergus octosetaceus* mitogenome.

Small differences were found when comparing with a museum mitogenome, derived from an Argentinian individual collected in the 1950’s [15]. The museum specimen assembly is 452 bp smaller and composed of 70.37% coding regions, and it has an incomplete D-loop sequence and a truncated OH subunit.

### Synteny conservation

Chromosomal collinearity analysis revealed extensive syntenic conservation across the Anatidae family (**Fig. 4**). Multiple inversion events were identified, particularly on chromosomes 1, 2, and 3. Microchromosomes and the Z chromosome showed strong structural conservation, with syntenic blocks often spanning their entire lengths. All evaluated Anatidae species share the same karyotype (2n = 40). However, *O. jamaicensis* exhibits a derived fused chromosome 4 (resulting from the fusion of chromosome 4 and a microchromosome), a feature also found in *Gallus gallus*.

**Figure 4.**
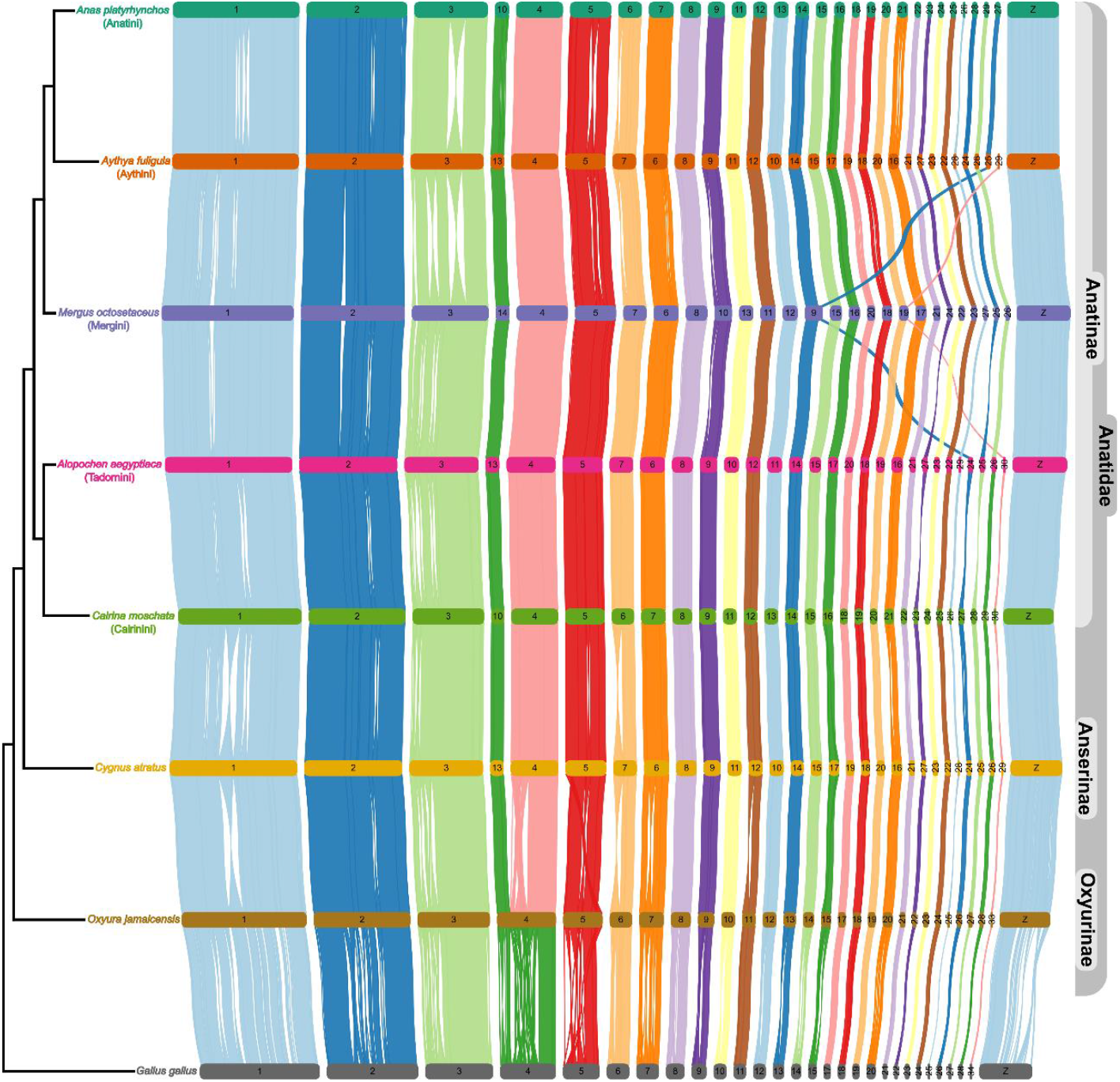
Structural synteny between chromosomes from Anatidae family species with *Gallus gallus* as an outgroup. The topology is based on Buckner et al. [49].

### Demographic history

The demographic history analysis with PSMC revealed dynamic fluctuations in the historical effective population size (N_e_) of the Brazilian merganser (**Fig. 5**). N_e_ declined gradually until a pronounced reduction around ~150 thousand years ago (kya). This decline occurred across the glacial cycles of the Mid-Pleistocene until the Eemian interglacial period (130–115 kya). Later, N_e_ rebounded and peaked between 40–30 kya, while temperatures gradually reduced. After 30 kya, temperatures dropped further to their lowest value during the Last Glacial Maximum (LGM; ~26–19 kya), when Ne began a more pronounced and relatively ‘rapid’ decrease. The effective population size (N_e_) did not recover from this decline until the inference limit at 10 kya.

**Figure 5.**
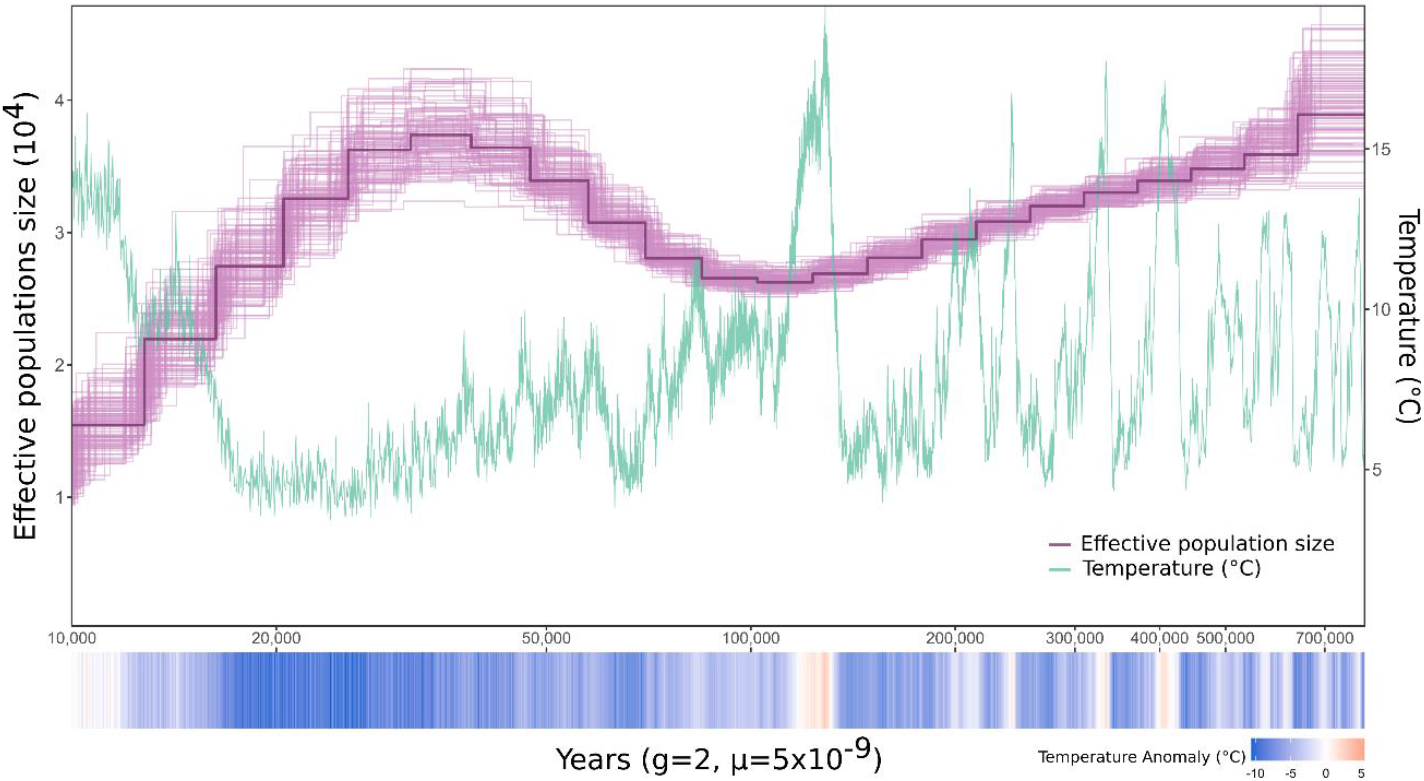
Demographic history estimated by Pairwise Sequentially Markovian Coalescent – PSMC. Estimated effective population size (N_e_) through time is depicted by the purple line, while the green line indicates global temperature variation measured in degrees Celsius (ºC). The barcode plot in the x axis indicates the mean temperature anomaly over time, relative to the 1950’s average global temperature of 14

## Discussion

We assembled the first chromosome-level genome of *M. octosetaceus*, achieving high completeness and minimal gaps. The genome size is consistent with expectations for birds and exhibits a high N50 (83.8 Mb), comparable to other recent chromosome-level assemblies in the Anatidae family. We identified 36 autosomal chromosomes and both sex chromosomes (Z and W), closely matching the expected karyotype for the group (n = 40). As it is characteristic of birds, several microchromosomes were present, which poses challenges for assembly and curation.

While analyzing the genomic composition, we observed an elevated proportion of repetitive elements when compared to other birds, which appears to be the result of a recent expansion of transposable elements. This is supported by a distinct peak (K2P=0) in **Fig. 2B**, reflecting the recent expansion of newly active repetitive elements. Furthermore, we observed a higher concentration of repetitive elements in macrochromosomes than in microchromosomes. This pattern is consistent with previous findings [50,51], suggesting that microchromosome gene enrichment results from a gradual loss of repetitive regions.

In addition to the nuclear genome, we also compared our newly assembled mitogenome to a previously published annotated assembly of an old specimen of museum from a likely extinct population of Argentina (**Supplementary Fig. S3**). Our version of the *M. octosetaceus* mitogenome is longer and does not present fragmented or incomplete regions. This difference is mainly because the former assembly was based on an old museum sample, which affects the quality and integrity of the extracted DNA, and posterior sequencing. In contrast, our fresh sample allowed us to properly reconstruct the complete mitogenome with all its expected genes.

In addition to those findings, the synteny analysis of this assembly provided insights into the particularities of chromosomal evolution within the Anatidae. Besides an overall syntenic conservation in the group, several inversion events were observed, and the apparent bias towards macrochromosomes is likely due to their higher proportion of repetitive elements. While the chromosome 4 (chr4) fusion in *O. jamaicensis* (unlike in *G. gallus*) retains 2n = 80 [52], Beklemisheva et al. [53] report similar rearrangements in Anserinae (fused/split chr4; 2n=80). Collectively, this suggests that karyotype conservation in Anatidae occurs through structural changes.

PSMC analysis suggests that fluctuations in N_e_ overlapped with major climatic events. During the Middle Pleistocene, successive glaciations produced significant temperature instability, which appears to have undermined N_e_ (**Fig 5**). Following the warmer Eemian interglacial period, temperatures stabilized within a cooler range of approximately 5-10°C. This period of relative climatic stability, represented by the bluer shading in **Fig. 5**, could allow the recovery of Ne. However, this trend reversed with the onset of the severe glacial conditions of the LGM, leading to a more pronounced reduction in N_e_. Thus, our findings suggest that temperature may have been a significant driver of these fluctuations. It is particularly concerning, considering the critical situation of the Brazilian mergansers that are restricted to few remaining populations in the wild, and a recent modelling distribution study [54] indicates that other suitable areas are outside of protected areas, and under severe threat due to construction of hydroelectric plants.

## Conclusion

This study provides a high-quality chromosome-level genome for *M. octosetaceus*, including a complete and intact mitogenome. Repetitive element analysis raises questions about which specific elements are expanding and their potential impacts on genome evolution in birds. Comparative genomic studies with other Anatidae species are now needed to elucidate chromosomal evolution, for which this genome will be a fundamental resource.

The demographic history reconstruction further emphasizes the species’ vulnerability, revealing past contractions in N_e_ associated to climate changes that likely contributed to reduced contemporary genetic diversity and persistent threats to natural populations. However, the drivers underlying these N_e_ fluctuations remain unresolved.

While temperature appears to be a contributing factor, comprehensive niche modeling studies incorporating multiple variables are essential to elucidate this pattern.

Overall, this reference genome addresses key questions and will serve as a vital resource for future research to understand the remnant populations dynamics, devise in-situ management strategies, and eventually support reintroduction efforts for this critically endangered species.

## Data Availability

The *Mergus octosetaceus* (Brazilian merganser) assemblies are available on NCBI under accession number GCA_036873955.1 (BioProject PRJNA1072166) for haplotype 1, and GCA_036850655.1 (BioProject PRJNA1072165) for haplotype 2. All PacBio and Hi-C raw data are available and have been deposited on the Sequence Read Archive (SRA) from NCBI (SRS22164953).

## Abbreviations

BUSCO: Benchmarking Universal Single Copy Orthologs
CDS: Protein-coding genes
KYA: Thousand years ago
N_e_: Effective population size
OH: Replication origin
PSMC: Pairwise Sequentially Markovian Coalescent
rRNA: Ribosomal RNA
tRNA: Transfer RNA
VGP: Vertebrate Genomes Project.

## Acknowledgements

We thank Larissa S. Arantes for valuable insights; Diego De Panis for providing his scripts from the jATG project (https://github.com/diegomics/jATG/) as a source of inspiration, and Léo Marujo for the illustration of the Brazilian merganser.

## Author Contributions

HPGN: Writing - Original Draft Preparation, Investigation, Formal Analysis; IRFB: Investigation (Repetitive/Synteny); DBC: Investigation (Mitogenome); DPC: Sample collection; KH: Sample coordinator; JB: Wet lab director; BO: Hi-C sequencing; TT: Hi-C sequencing; NJ: Pacbio sequencing; LA: Genome assembly; NB: Genome curation; GF: Bioinformatics lead; OF: Genome sequencing coordinator; EDJ: Genome sequencing coordinator; FRS: Conceptualization, Funding acquisition, Supervision, Writing - Review & Editing. All authors read and approved the manuscript.

## Supplementary Materials

**Supplementary Fig S1.**
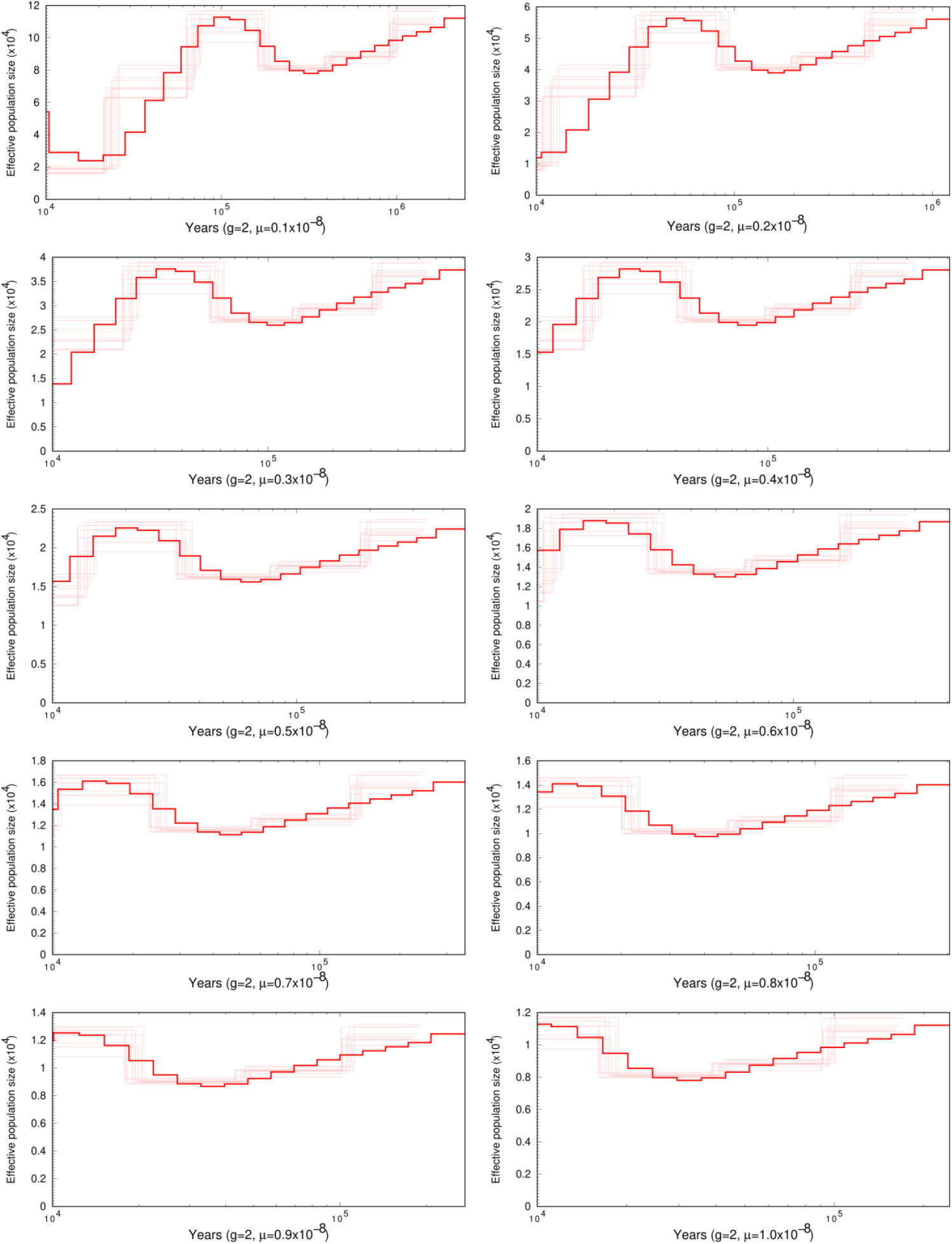
Pairwise Sequentially Markovian Coalescent (PSMC) mutation rate tests with fixed parameters (N30 t15 r4 p4+5X3+4). We used the *Gallus gallus* mutation rate interval from Bergeron et al. (2023).

**Supplementary Fig. S2.**
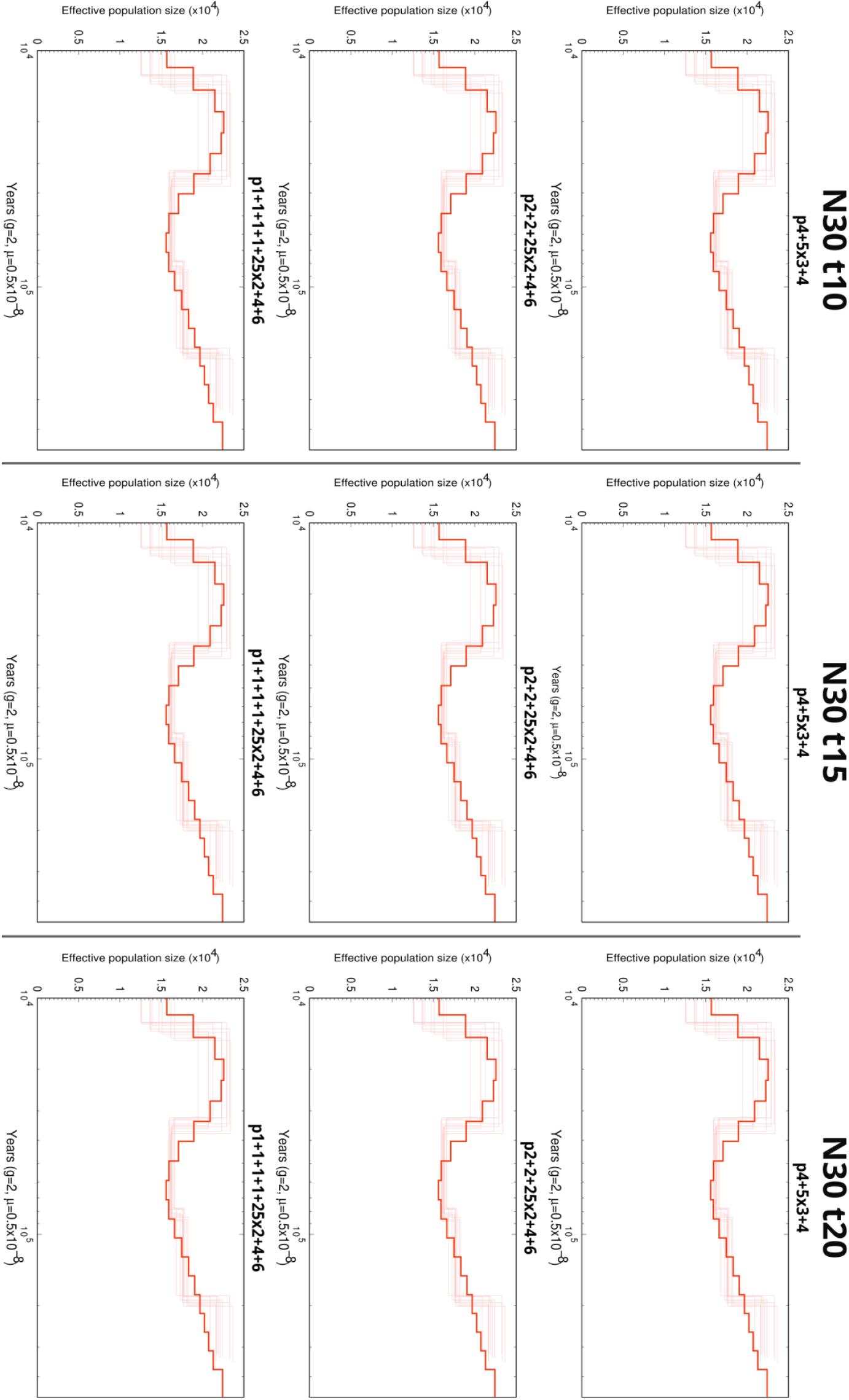
Pairwise Sequentially Markovian Coalescent (PSMC) maximum coalescent and pattern tests with a fixed mutation rate (5×10^−9^).

**Supplementary Fig. S3.**
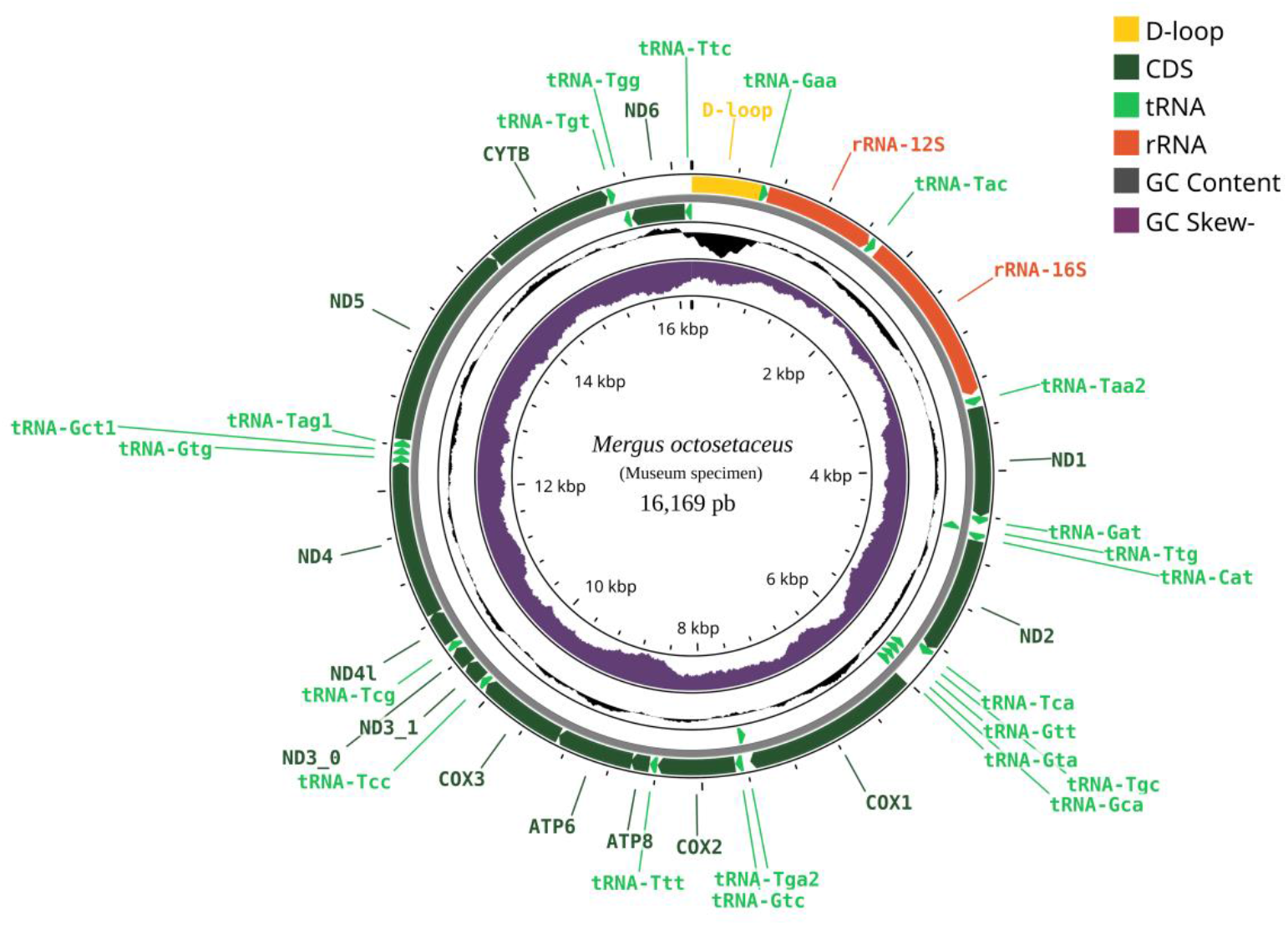
Structural annotation of the museum specimen of *Mergus octosetaceus* mitogenome.

**Table S1.**
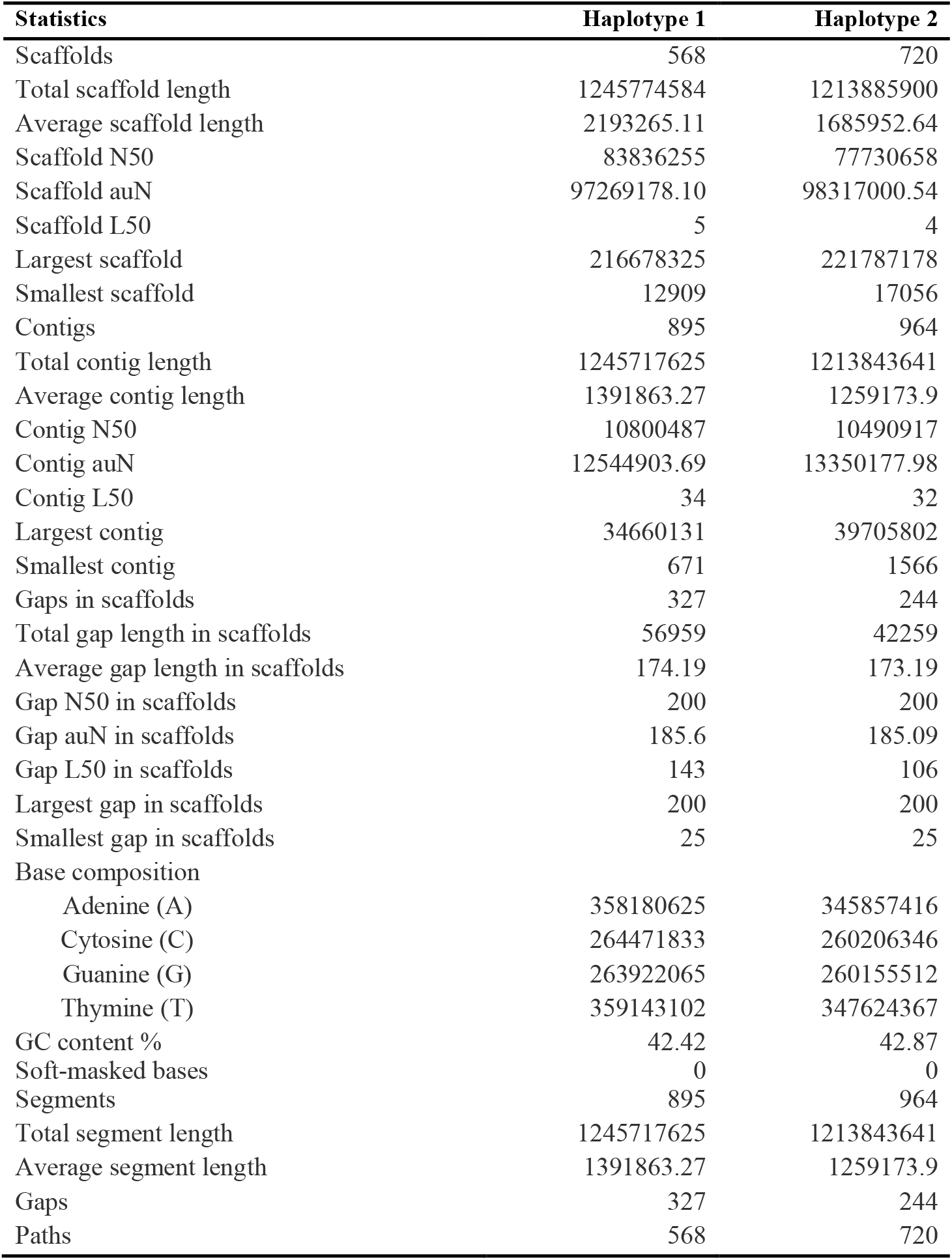
Primary assembly statistics from both haplotypes.

| **Statistics** | **Haplotype 1** | **Haplotype 2** |
| --- | --- | --- |
| Scaffolds | 568 | 720 |
| Total scaffold length | 1245774584 | 1213885900 |
| Average scaffold length | 2193265.11 | 1685952.64 |
| Scaffold N50 | 83836255 | 77730658 |
| Scaffold auN | 97269178.10 | 98317000.54 |
| Scaffold L50 | 5 | 4 |
| Largest scaffold | 216678325 | 221787178 |
| Smallest scaffold | 12909 | 17056 |
| Contigs | 895 | 964 |
| Total contig length | 1245717625 | 1213843641 |
| Average contig length | 1391863.27 | 1259173.9 |
| Contig N50 | 10800487 | 10490917 |
| Contig auN | 12544903.69 | 13350177.98 |
| Contig L50 | 34 | 32 |
| Largest contig | 34660131 | 39705802 |
| Smallest contig | 671 | 1566 |
| Gaps in scaffolds | 327 | 244 |
| Total gap length in scaffolds | 56959 | 42259 |
| Average gap length in scaffolds | 174.19 | 173.19 |
| Gap N50 in scaffolds | 200 | 200 |
| Gap auN in scaffolds | 185.6 | 185.09 |
| Gap L50 in scaffolds | 143 | 106 |
| Largest gap in scaffolds | 200 | 200 |
| Smallest gap in scaffolds | 25 | 25 |
| Base composition |  |  |
| Adenine (A) | 358180625 | 345857416 |
| Cytosine (C) | 264471833 | 260206346 |
| Guanine (G) | 263922065 | 260155512 |
| Thymine (T) | 359143102 | 347624367 |
| GC content % | 42.42 | 42.87 |
| Soft-masked bases | 0 | 0 |
| Segments | 895 | 964 |
| Total segment length | 1245717625 | 1213843641 |
| Average segment length | 1391863.27 | 1259173.9 |
| Gaps | 327 | 244 |
| Paths | 568 | 720 |

**Table S2.**
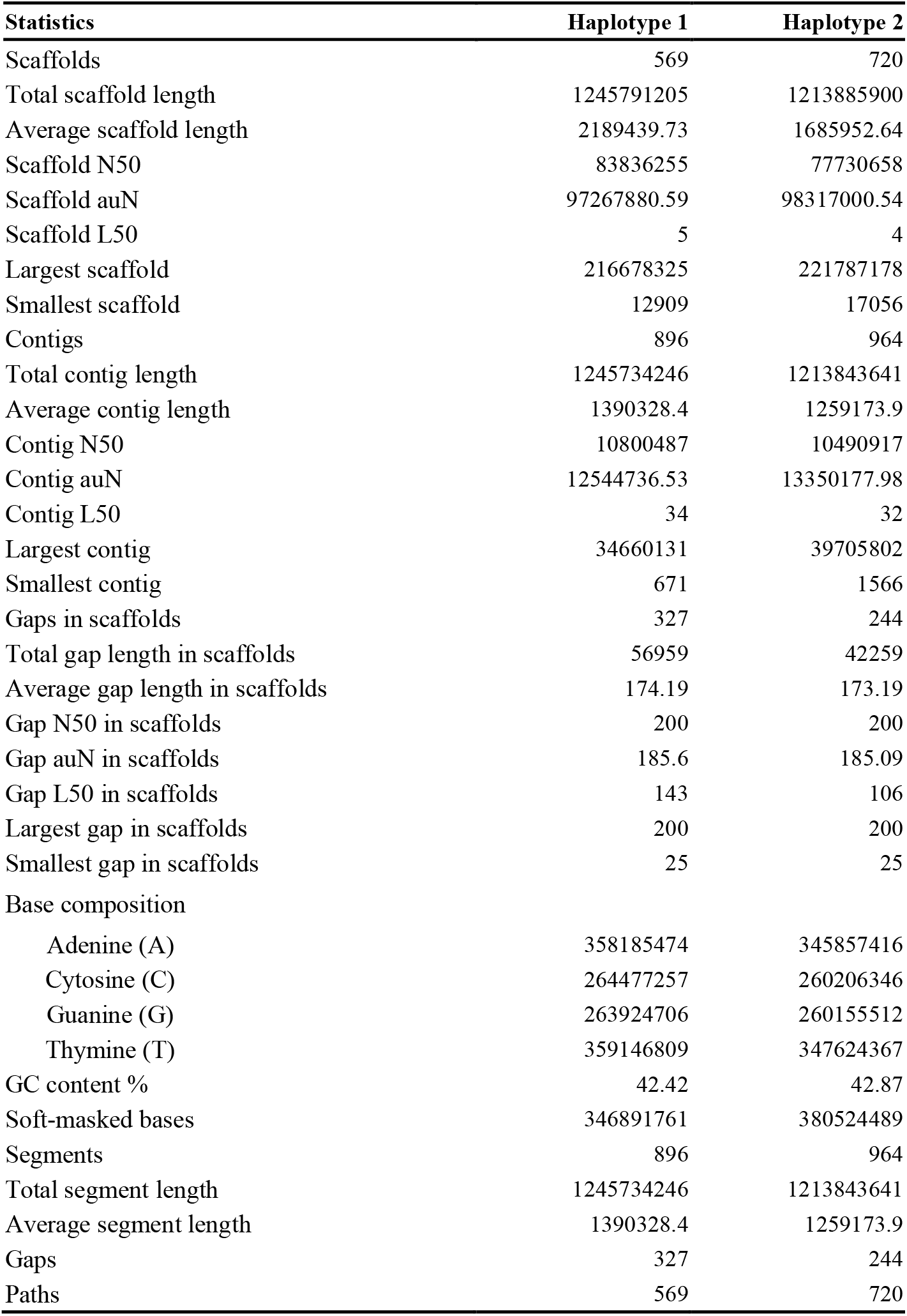
Curated assembly statistics from both haplotypes.

